# Constrained evolutionary paths to macrolide resistance in a *Neisseria* commensal converge on ribosomal genes through sequence duplication

**DOI:** 10.1101/2021.04.05.438469

**Authors:** Jordan C. Raisman, Michael A. Fiore, Lucille Tomin, Joseph K.O. Adjei, Virginia Aswad, Jonathan Chu, Christina J. Domondon, Ben A. Donahue, Claudia A. Masciotti, Connor G. McGrath, Jo Melita, Paul A. Podbielski, Madelyn R. Schreiner, Lauren J. Trumpore, Peter C. Wengert, Emalee A. Wrightstone, André O. Hudson, Crista B. Wadsworth

## Abstract

*Neisseria* commensals are an indisputable source of resistance for their pathogenic relatives; however, the evolutionary paths commensal species take to reduced susceptibility in this genus have been relatively underexplored. Here, we leverage *in vitro* selection as a powerful screen to identify the genetic adaptations that produce azithromycin resistance (≤ 2 μg/mL) in the *Neisseria* commensal, *N. elongata*. Across multiple lineages (n=7/16), we find mutations encoding resistance converge on the gene encoding the 50S ribosomal L34 protein (*rpmH*) and the intergenic region proximal to the 30S ribosomal S3 protein (*rpsC)* through duplication events. Importantly, one of the laboratory evolved mutations in *rpmH* is identical, and two nearly identical, to those recently reported to confer high-level resistance to azithromycin in *N. gonorrhoeae*. Transformations into the ancestral *N. elongata* lineage confirmed the causality of both *rpmH* and *rpsC* mutations. Though most lineages inheriting duplications suffered *in vitro* fitness costs, one variant showed no growth defect, suggesting the possibility that it may be sustained in natural populations. Finally, we assessed the potential of horizontal transfer of derived resistance mutations into multiple strains of *N. gonorrhoeae*. Though we were unable to transform *N. gonorrhoeae* in this case, studies like this will be critical for predicting commensal alleles that are at risk of rapid dissemination into pathogen populations.

**Importance:** Commensal bacterial populations have been increasingly recognized for their importance as sources of resistance for pathogens, however the collection of antimicrobial resistance (AMR) mechanisms within these communities are often understudied. The risk of reduced antibiotic susceptibility as a result of horizontal gene transfer (HGT) is amplified in highly recombinogenic genera, such as the *Neisseria*. Indeed, there have been multiple documented cases of macrolide and beta-lactam resistance acquisition in the pathogen *N. gonorrhoeae* from close commensal relatives. This work uncovers multiple novel azithromycin resistance-conferring mutations in a *Neisseria* commensal through experimental evolution, investigates their fitness impacts, and explores the possibility of transfer to *N. gonorrhoeae*. Ultimately these types of studies will illuminate those resistance mutations that may rapidly be acquired across species boundaries.

## Introduction

Commensal bacterial populations have been increasingly recognized for their importance as sources of adaptive genetic variation for pathogens through horizontal gene transfer (HGT). This threat of rapid evolution as a result of DNA donation is especially amplified in highly recombinogenic genera, such as the *Neisseria*. Members of this genus readily donate DNA to one another through pilus-mediated *Neisseria*-specific DNA uptake and homologous recombination (1-3). Observations of genetic mosaicism, whereby loci within a particular lineage have been acquired from another species, are common (4-11) and occur genome-wide (12, 13). This promiscuous allelic exchange has been documented to have facilitated rapid adaptive evolution of important phenotypic characteristics such as antimicrobial resistance (6, 8-10) and body-site colonization niche shifts (11); and overall, recently incorporated mosaic sequences show signatures consistent with positive selection (12), suggesting that intragenus recombination is an important source of beneficial genetic variation.

The genus *Neisseria* is comprised of several Gram-negative species that typically colonize the mucosa of humans and animals. Most human-associated *Neisseria* inhabit the naso- and oropharynx, and are carried harmlessly as commensals in 10-15% of healthy human adults and children (14, 15). Only one species, *N. gonorrhoeae* (Ngo), is an obligate pathogen which also colonizes the urogenital tract and rectum, and causes the sexually transmitted infection gonorrhea (16). In recent years it has become increasingly clear that *Neisseria* commensals serve as important reservoirs of adaptive genetic variation for Ngo through HGT (6, 8, 9, 12, 17); however, most of the non-pathogenic *Neisseria* have been infrequently characterized as they rarely cause systemic or life-threatening disease, except in immunocompromised individuals (18, 19).

Horizontal transfer and subsequent spread of commensal *Neisseria* resistance mechanisms has historically played a large role in rendering antibiotic therapies ineffective in Ngo. Reduced drug susceptibility in commensal *Neisseria* populations has been shown to be directly selected for after antibiotic usage (20), and thus these species will always be a persistent threat for resistance donation. In a U.S. gonococcal population dataset, over 11% of reduced susceptibility to azithromycin was acquired through inheritance of commensal *transferable efflux pump* (*mtr*) alleles (8-10). Additionally, the majority of resistance to third-generation cephalosporins in gonococci is derived through mosaic *penicillin-binding protein 2* (*penA*) alleles gained from close commensal relatives (6, 10). Though studies that characterize the resistance genotypes and phenotypes in panels of commensal *Neisseria* (such as (17, 21, 22)) will aid in prospective surveillance for novel resistance that may be rapidly acquired by Ngo, the utility of these approaches is ultimately limited by extensive under sampling of natural commensal populations.

Experimental evolution can be leveraged as an alternative to phenotyping and genotyping large panels of bacteria, as a powerful screen for the genetic adaptations that underly antibiotic resistance in the commensal *Neisseria*, and thus may be at risk of HGT to Ngo. Laboratory evolution experiments reveal the spontaneous mutations caused by DNA replication and repair errors which increase mean fitness in new selective environments (23), and are ideal to illuminate the mechanisms of adaptation in bacteria due to their short generation times. *In vitro* selection has previously been used successfully to nominate mechanisms underlying ceftriaxone and azithromycin reduced susceptibility in Ngo (24, 25), and thus is a promising tool for illuminating new mechanisms of resistance in *Neisseria* commensals.

In this study, we use experimental evolution to investigate the potential for a *Neisseria* commensal (*N. elongata*, Nel) to evolve resistance to the macrolide antibiotic azithromycin, which until this year (26) was recommended as a first-line antibiotic, in combination with ceftriaxone, for the treatment of uncomplicated cases of gonorrhea in the United States. Here, we characterize the evolutionary response of Nel replicate lineages to azithromycin and consider if there is a single or multiple adaptive solutions to selection. Furthermore, we assess the fitness costs of derived mutations, and their potential for transfer into Ngo.

## Results

### Evolutionary trajectories to macrolide resistance in N. elongata

*N. elongata* AR Bank #0945 was selected to explore the evolutionary paths to azithromycin resistance in a *Neisseria* commensal. *N. elongata* AR Bank #0945 has been tested for its minimum inhibitory concentration (MIC) to azithromycin (0.38 μg/mL) and sequenced (SAMN15454046) by our group previously (21). In this study, a draft genome was assembled as an ancestral reference for all derived lineages. The assembly contained 2,572,594 bps, and 2,509 annotated genes (JAFEUH000000000).

Selective pressure was applied to sixteen replicates of AR Bank #0945 by creating a concentration gradient of azithromycin on standard growth media via application of Etest strips (Figure 1A). Cells were selected for passaging by sweeping the entire zone of inhibition (ZOI) and a 1 cm band in the bacterial lawn adjacent to the ZOI (Figure 1A). After 20 days, or approximately 480 generations, the average MIC value for all evolved cell populations increased to 7.6 μg/mL, which was significantly higher compared to day one values (*W* = 98.5, *P* = 0.0004), and ranged from 0.19 to 48 μg/mL (Figure 1B and C). Control populations with no drug selection (n=4), showed no increase in MIC compared to the ancestral stock. To mitigate the possibility of heterogenous cultures at the termination of the experiment, a single colony was selected from each evolved and control population for further MIC testing and genomic sequencing. MIC values for drug-selected single colony picks tended to be higher than those recorded for population values, and ranged from 0.5 to 64 μg/mL, with an average value of 14.4 μg/mL (Table 1).

**Table 1.** Azithromycin MICs and derivation data for the strains used in this study.

| Strain | Azi MIC (µg/ml) | Recipient strain | Donor Strain |
| --- | --- | --- | --- |
| **Azithromycin selected strains t** |  |  |  |
| AM1 | 24 | - | - |
| CBW1 | 1 | - | - |
| CBW2 | 1 | - | - |
| CBW3 | 1 | - | - |
| CBW4 | 64 | - | - |
| CBW5 | 0.5 | - | - |
| CBW6 | 64 | - | - |
| EW1 | 0.5 | - | - |
| JA1 | 8 | - | - |
| JC1 | 0.5 | - | - |
| JM1 | 0.75 | - | - |
| JR1 | 1 | - | - |
| LJT1 | 16 | - | - |
| LT1 | 24 | - | - |
| MRS1 | 24 | - | - |
| PAP1 | 0.75 | - | - |
| **Control strains, no drug selection** |  |  |  |
| ND1 | 0.38 | - | - |
| ND2 | 0.38 | - | - |
| ND3 | 0.38 | - | - |
| ND4 | 0.38 | - | - |
| **Transformants b** |  |  |  |
| T-CBW4-1 | 48 | AR-0945 | CBW4 |
| T-CBW4-2 | 48 | AR-0945 | CBW4 |
| T-CBW4-3 | 48 | AR-0945 | CBW4 |
| T-CBW6-1 | 64 | AR-0945 | CBW6 |
| T-CBW6-2 | 64 | AR-0945 | CBW6 |
| T-CBW6-3 | 64 | AR-0945 | CBW6 |
| T-LT1-1 | 24 | AR-0945 | LT1 |
| T-LT1-2 | 24 | AR-0945 | LT1 |
| T-LT1-3 | 24 | AR-0945 | LT1 |
| T-MRS1-1 | 24 | AR-0945 | MRS1 |
| T-MRS1-2 | 24 | AR-0945 | MRS1 |
| T-MRS1-3 | 24 | AR-0945 | MRS1 |
| T-JA1-1 | 8 | AR-0945 | JA1 |
| T-JA1-2 | 8 | AR-0945 | JA1 |
| T-JA1-3 | 8 | AR-0945 | JA1 |
| T-LJT1-1 | 12 | AR-0945 | LJT1 |
| T-LJT1-2 | 12 | AR-0945 | LJT1 |
| T-LJT1-3 | 12 | AR-0945 | LJT1 |
| T-AM1-1 | 24 | AR-0945 | AM1 |
| T-AM1-2 | 24 | AR-0945 | AM1 |
| T-AM1-3 | 12 | AR-0945 | AM1 |

t. Selection on an azithromycin concentration gradient; b. Selection on 4 µg/ml azithromycin

**Figure 1.**
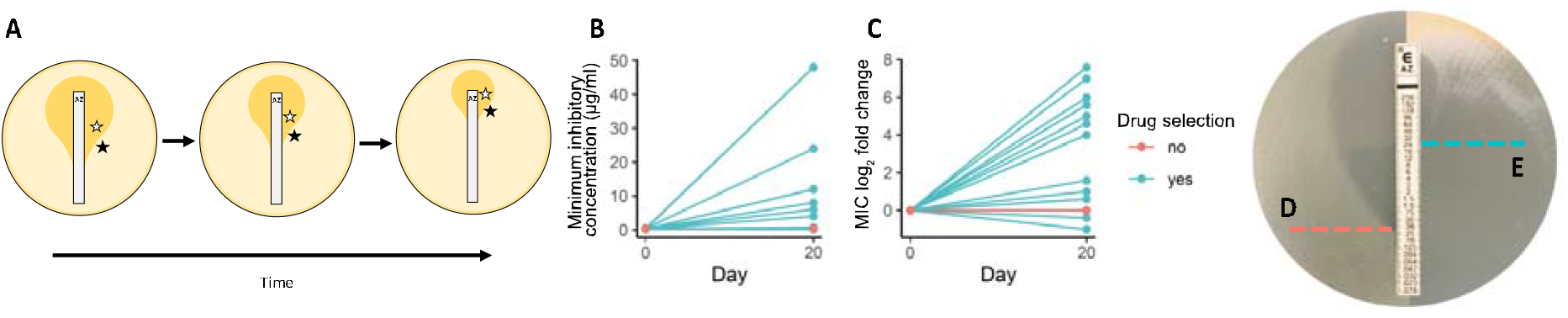
*In vitro* evolution under azithromycin selection produces high-level resistance in multiple replicate lineages of the *Neisseria* commensal, *N. elongata*. (A) Parallel cultures of *N. elongata* AR Bank #0945 were passaged across an Etest-generated concentration gradient of azithromycin, selecting any cell growth in the zones of inhibition (white stars) and a 1 cm band at the edge of the bacterial lawn (black stars) for 20 days (n=16 replicates). (B,C) In 7 of the 16 lineages, reduced susceptibility as defined by the CLSI cutoff of ≥ 2 μg/mL emerged. In all cases, MICs for evolved lineages increased from the ancestral *N. elongata* AR Bank #0945 strain value of 0.38 μg/mL (D), with strain LT1 displayed (E).

Evolved cell lines were sequenced and aligned to the *N. elongata* AR Bank #0945 draft assembly to nominate derived polymorphisms. Mutations that were shared with control strains, or those shared with ancestral reads mapped back to the reference assembly, were not further considered. In total, 37 derived mutations were identified across all sequenced strains (Table 2). The most frequently observed mutations were found in the glucokinase encoded by *glk* (n=13 lineages), followed by those in *rpmH* encoding the 50S ribosomal protein L34 (n=4), and mutations in the intergenic region proximal to *rpsC* encoding the 30S ribosomal S3 protein (n=3). Unique mutations observed within annotated genes, were present in the coding domains for: the capsular polysaccharide phosphotransferase (encoded by *cps12A*), isocitrate lyase (*aceA*), RNA 2’-phosphotransferase (*kptA*), the di-/tripeptide transporter (*dptT*), and the bifunctional (p)ppGpp synthase/hydrolase (*spoT*). All strains with azithromycin MIC values ≥ 2 μg/mL (the Clinical & Laboratory Standards Institute (CLSI) reduced susceptibility breakpoint for *N. gonorrhoeae* (27)) were associated inheritance of mutations in either *rpmH* or the intergenic region proximal to *rpsC*.

**Table 2.**
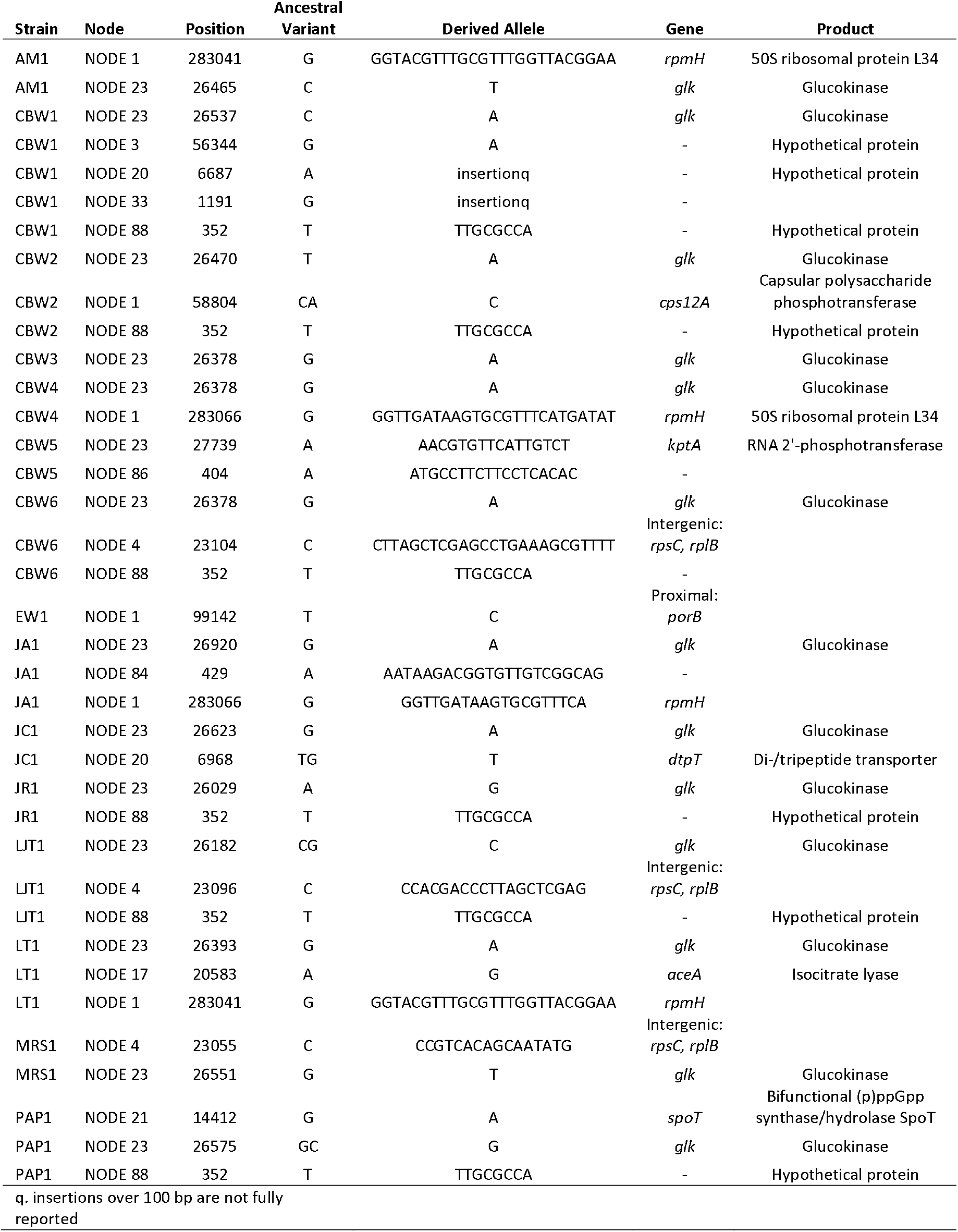
Derived SNPs in evolved lineages annotated in reference to the draft AR-0945 assembly (JAFEUH000000000)

### Confirmation of the causality of high-level resistance encoding mutations

To assess the causality of the mutations encoding macrolide resistance in *N. elongata* AR Bank #0945, the ancestral stock was transformed with genomic DNA from evolved cell lines with MIC values ≥ 2 μg/mL (n=7). Genomic DNA from evolved lineages successfully transformed the ancestral stock in all cases. To identify causal loci, three colonies from each transformation were selected to characterize the polymorphisms which had been inherited from donor strains that were not present in the AR Bank #0945 ancestral recipient. The only region inherited across replicate transformant colony picks contained either mutations in *rpmH* (DNA donated from AM1, CBW4, JA1, or LT1), or the intergenic region near *rpsC* (DNA donated from CBW6, LJT1, or MRS1) (Table 2). Sanger sequencing confirmed the identity and presence of these mutations (Figure 2), and translation of *rpmH* duplications at the amino acid level indicated the in-frame insertions: 8SVTKRKRT15, 7LKRTYQ12, and 8HIMKRTYQ15 (Figure 3).

**Figure 2.**
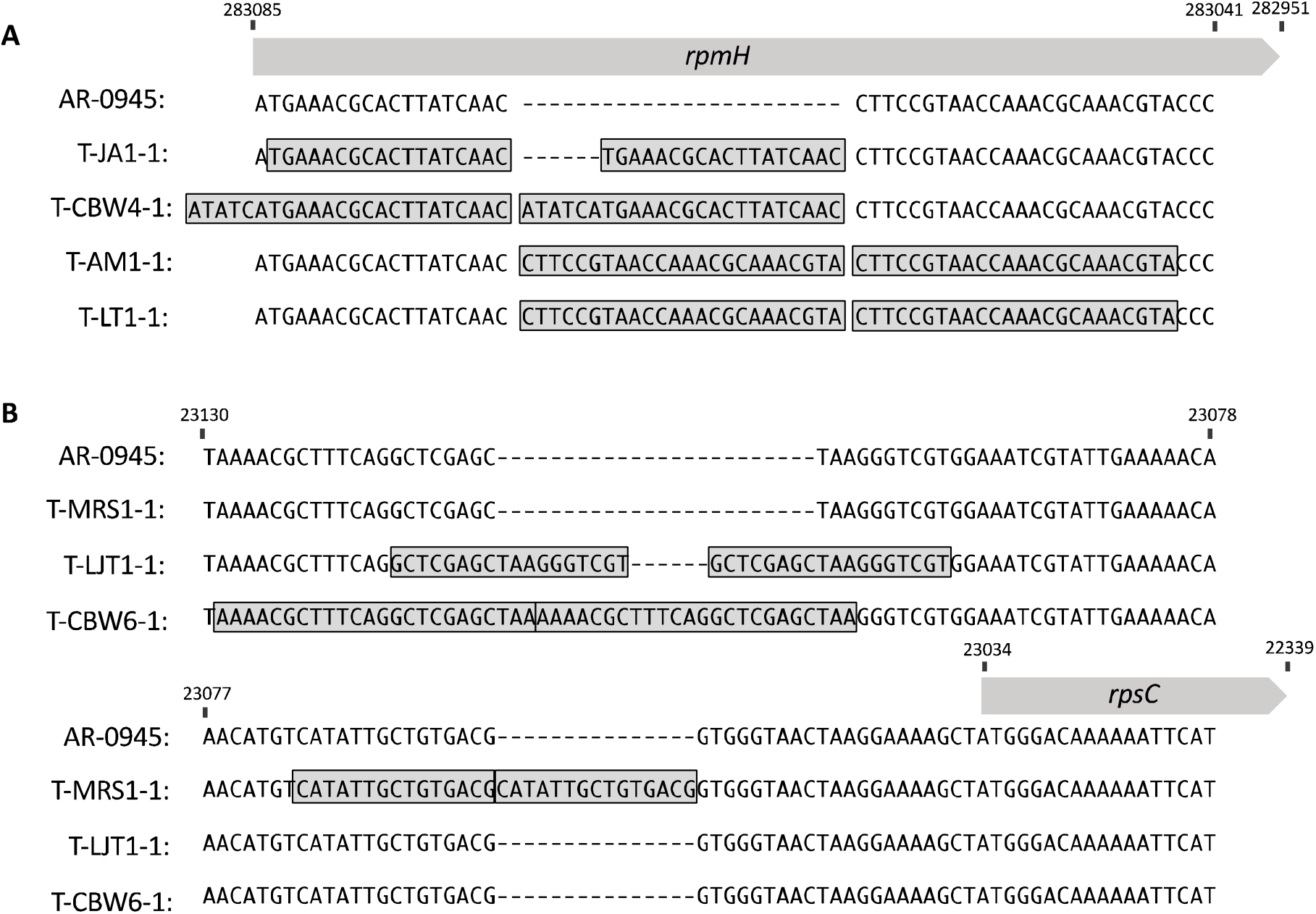
Repeated emergence of mutations within and proximal to genes encoding ribosomal proteins are causal to high-level azithromycin resistance. (A) Whole genomic sequencing revealed that four strains evolved duplications in *rpmH*, the gene encoding the 50S ribosomal protein L34. Transformants (T-AM1, T-CBW4, T-JA1, and T-LT1) generated from these lineages were Sanger sequenced to confirm the location and identity of nominated derived insertional polymorphisms, data shown. (B) The remaining mutations underlying high-level resistance (n=3) were located proximally to *rpsC*, which encodes the 30S ribosomal protein S3, and were also duplication events. Sanger sequences for the transformants (T-CBW6, T-LJT1, and T-MRS1) are displayed. Reference positions in Node 1 and Node 4 of the AR-0945 draft assembly are displayed for *rpmH* and *rpsC* respectively.

**Figure 3.**
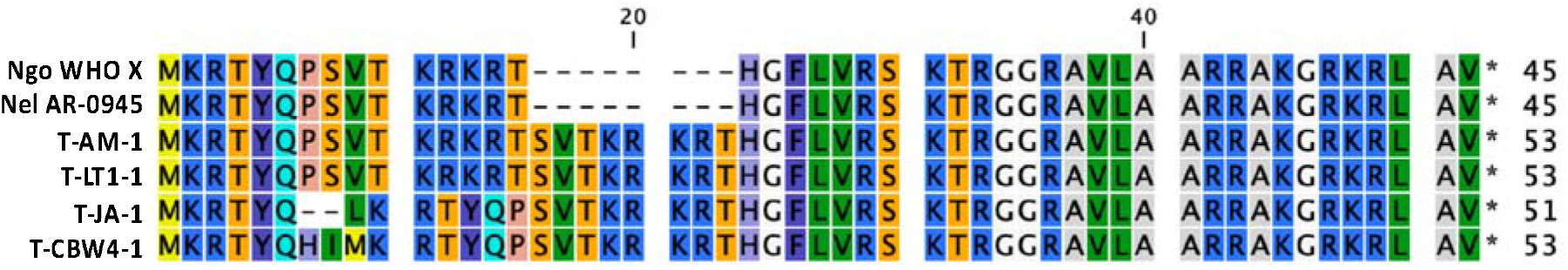
At both the nucleotide and amino acid level Nel AR-0945 RpmH is identical to the Ngo WHO F reference (<u>LT591897.1</u>) sequence. Evolved in-frame sequence duplications in *Nel* lineages introduce 8SVTKRKRT15, 7LKRTYQ12, and 8HIMKRTYQ15 insertions, which are identical to (7LKRTYQ12) and nearly identical to those recently discovered to confer high-level macrolide resistance in Ngo (25).

In most cases recovered transformants had azithromycin MICs that perfectly mirrored the donor strain phenotypes. However, all three transformants with CBW4 as a donor consistently had phenotypes of 48 μg/mL, one dilution below the donor strain phenotype of 64 μg/mL (Table 1). Additionally, LJT1 transformants had MIC values of 12 μg/mL, also one dilution below the donor strain phenotype (Table 1). Finally, one of the three AM1 transformants (T-AM1-3) had a lower MIC (12 μg/mL) than the other transformants and the AM1 donor strain (24 μg/mL).

### Most novel ribosomal variants reduce *in vitro* fitness

In order to evaluate the fitness costs of azithromycin resistance-conferring mutations, the optical densities of transformant cells lines were compared to the ancestral *N. elongata* AR Bank #0945 strain over a 21-hour period (Figure 4A). At hour 21, OD_600_ values for AR-0945 replicates ranged from 0.68 to 0.81 (n=6); and six of the seven transformants had significantly lower optical densities (Figure 4B; Tukey’s HSD, p < 0.001). OD_600_ values for T-LJT-1 however ranged from 0.70-0.78 (n=6), which were not significantly different compared to the ancestral strain (Figure 4B; Tukey’s HSD, p < 0.99).

**Figure 4.**
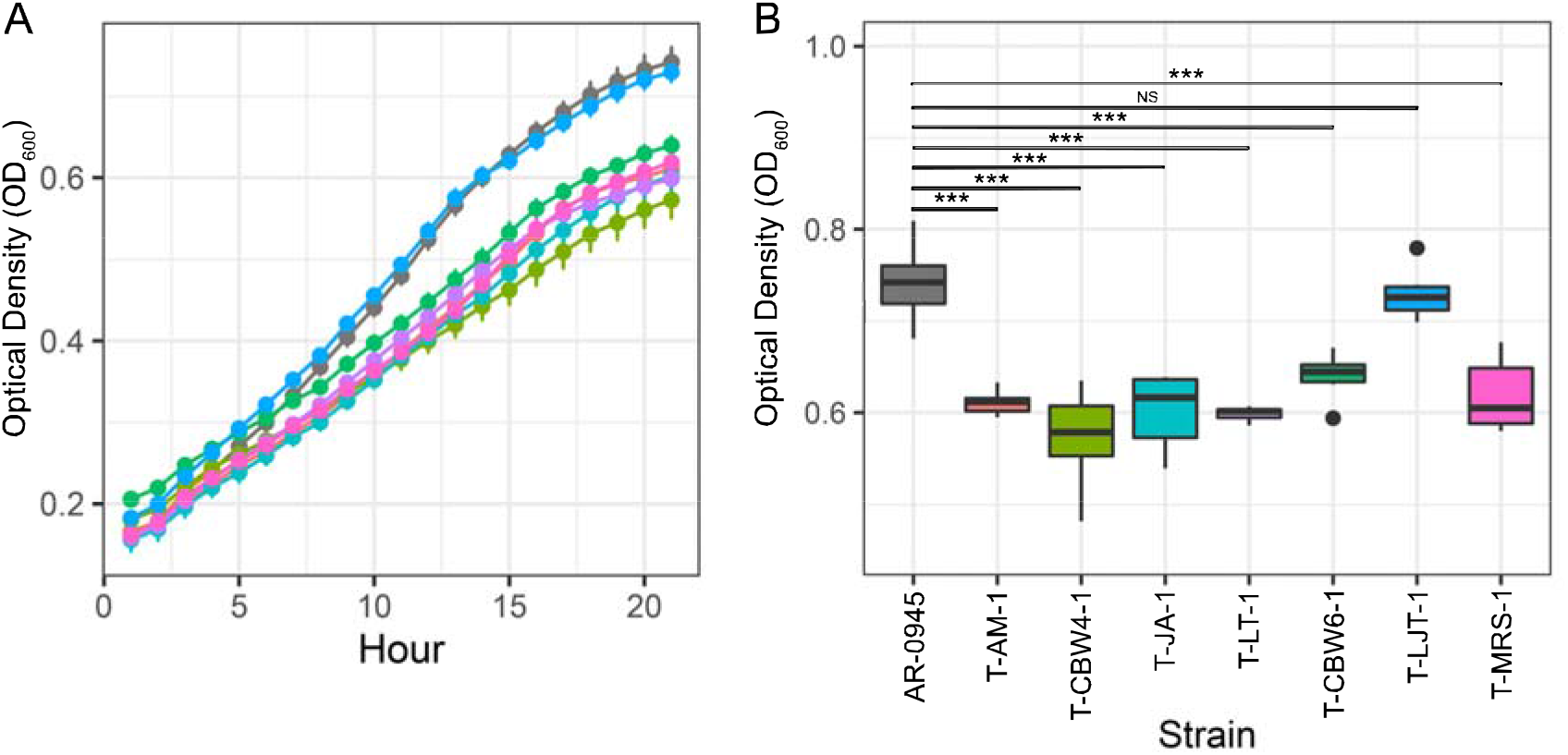
Impact of novel alleles in evolved cell lineages on growth rate compared to the ancestral AR-0945 strain. (A) Growth curves in GCP broth supplemented 1% Kelloggs solution were obtained by monitoring OD_600_ over 21 hours (n=6 replicates per strain). (B) Growth was significantly reduced in all strains compared to the ancestral at 21 hours, except for strain LJT. Significance between strains was determined using a one-way ANOVA followed post-hoc by Tukey’s HSD where ***□= *p* <□0.001 and NS = not significant.

### N. elongata ribosomal variants are inefficient or incapable of transfer to a pathogenic relative

*Neisseria* commensals are known sources of resistance for Ngo (6, 8-10), and therefore we test the ability of the evolved ribosomal duplication mutations uncovered in this study to be transferred into multiple Ngo strains. To establish a baseline rate of transformation for these alleles, we first quantify the number of transformants recovered for *N. elongata* AR Bank #0945 transformed with genomic DNA from evolved cell lines. In brief, we provided 100 ng of genomic DNA from evolved lineages in AR Bank #0945 cell suspensions, and after a 4-hour period allowing for the expression of any newly acquired alleles, selected on 4 μg/mL azithromycin. The percentage of colonies recovered on selective media as compared to colonies recovered on non-selective media was then calculated across three replicate trials, and averaged between 0.0003 to 0.0017 percent for all DNA (Figure 5).

**Figure 5.**
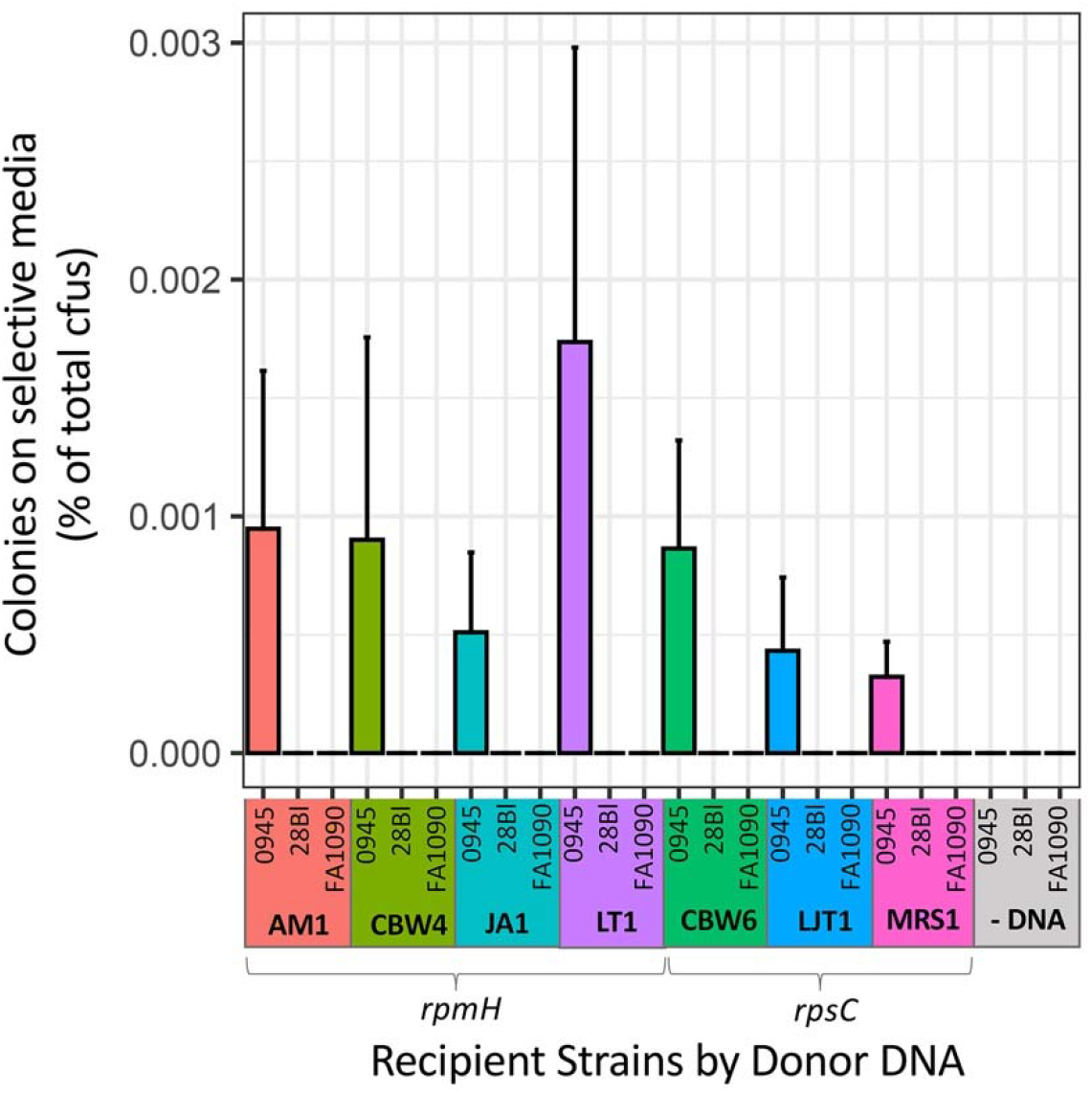
Genomic DNA from evolved resistant Nel lineages could transform the Nel AR-0945 ancestral strain to high-level macrolide resistance by inheritance of ribosomal duplication mutations. The colonies recovered on selective media (4 μg/mL azithromycin) as a percentage of total colony forming units (cfus) recovered on non-selective media ranged from 0.0003 to 0.0017 (n=3 trials per recipient strain and donor DNA combination). However, DNA containing Nel ribosomal variants were inefficient or incapable of transfer to naturally competent pilated Ngo 28Bl and FA1090 strains, with no colonies recovered. Additionally, no colonies were observed on selective media after transformation without template DNA for both Nel and Ngo reactions.

Though in natural populations of *Neisseria* HGT from commensals to Ngo has been repeatedly demonstrated, recent work has shown that commensal DNA is toxic to Ngo (28, 29). Ngo uptakes DNA through a Type IV pilus (Tfp)-based system, which binds preferentially to *Neisseria*-specific DNA (1-3). This DNA is then transported across the cell wall, and becomes integrated via homologous recombination into the genome by RecA. However, since Nel and Ngo have different intrinsic methylases, Ngo restriction enzyme(s) cleave incorporated DNA at heteroduplexes with Nel methylation signatures, resulting in the loss of chromosome integrity and cell death (28, 29). Nel DNA has been shown to be rendered less toxic through *in vitro* methylation using M.CviPI and M.SssI methyltransferases to modify cytosines in CpG and GpC motifs (28, 29).

Thus, transformation of Nel DNA was attempted with methylation modification (via M.CviPI and M.SssI methyltransferases) into naturally competent pilated Ngo 28Bl and FA1090 stocks. In all cases we recovered no colonies on selective media (Figure 5), following the same transformation protocol as used for Nel within this study, and which we have used previously for transformation of Ngo with Ngo DNA (8).

## Discussion

Multiple studies have demonstrated that commensal *Neisseria* serve as reservoirs of resistance for Ngo (6, 8-10), however a comprehensive evaluation of the resistance alleles commensals can harbor, though underway (17, 21, 22), and their likelihood of transfer to pathogenic relatives is still in its infancy. Here, we use experimental evolution to screen for the mutations that impart azithromycin resistance in Nel and assess their potential for transfer to Ngo.

Overall, our results support constrained evolutionary trajectories to high-level macrolide resistance in Nel. A diversity of azithromycin resistance mutations have been reported in Ngo, including: alterations in the 23S rRNA azithromycin binding sites (C2611T and A2059G) (10, 30, 31), a G70D mutation in the RplD 50S ribosomal protein L4 (32), *rplV* tandem duplications (10), variants of the rRNA methylase genes *ermC* and *ermB* (33), and mutations that enhance the expression of Mtr or increase the binding efficiency of MtrD with its substrates (8, 9, 34-36); however, high-level resistance is most frequently imparted by the aforementioned ribosomal mutations (10, 32). Similarly, for all cases of high-level resistance emergence in Nel drug-selected lineages within this study, causal mutations converged on either tandem duplications in the gene encoding the 50S ribosomal L34 protein (*rpmH*) or the intergenic region proximal to the 30S ribosomal S3 protein (*rpsC)*.

Fascinatingly, the evolved L34 mutations in Nel are identical or nearly identical to those recently reported in Ngo. High-level resistance in Ngo was found to be conferred by the L34 duplications 7PSVTKRKR14, 7PSVTNTYQP14, and 7LKRTYQ12 (25); and occurred in the same location as the Nel duplications 8SVTKRKRT15, 8HIMKRTYQ15, and 7LKRTYQ12 uncovered in this study (Figure 3). Preserved identity and location of *rpmH* mutations across *Neisseria* species suggests that this may be a conserved mechanism of rapid macrolide resistance acquisition across the genus. Though we do not find any of the other Ngo mutations in ribosomal components (C2611T, A2059G, *rplD* G70D, or *rplV* tandem duplications) in Nel, we describe novel azithromycin resistance conferring mutations in the intergenic region upstream of *rpsC*, similarly, produced through tandem duplication events (Figure 2). To our knowledge these have not yet been reported in *Neisseria*, and likely impact the expression of *rpsC* due to their location in or proximal to the promoter.

Almost all of the tandem duplications reported here imparted some fitness cost, as measured by *in vitro* growth assays of isogenic cell lines (Figure 4). Ngo duplication mutations in L34 were reported to be transitory and repeatedly lost in culture (25), further suggesting a fitness cost in alternate genetic backgrounds. However, the *rpsC* duplication in T-LJT-1 appeared to have no impact on fitness (Figure 4), suggesting that it may be sustained in natural populations. Further work will be needed to elucidate the long-term stability of all uncovered mutations, and to assess if they are either transitory stepping-stones or persistent, when coupled with compensatory mutations (e.g., as is the case for a variant in *acnB* mitigating growth defects in ceftriaxone resistant *penA* mutants (37)), mechanisms of macrolide resistance in commensal *Neisseria* species.

Assessing the likelihood of commensal resistance alleles to be transferred to Ngo will aid in determining those resistance mechanisms most at risk of rapid dissemination into pathogen populations; and may guide future prospective genotyping practices during routine public health surveillance efforts. Here, we attempted to transform novel Nel alleles into Ngo genetic backgrounds, however we were unable to recover transformants in all cases (Figure 5). One explanation for failed transformations could be insufficient sequence homology for RecA-mediated homologous recombination. Though, this is unlikely as *rpmH* shares 87% nucleotide and 100% amino acid identity between *N. elongata* AR Bank #0945 and to both Ngo FA1090 and 28Bl; similar to the percent identity of mosaic *mtr* sequences (∼90%) which we were previously able to transform into the 28Bl background (8). The sequence region including *rpsC* and 200 bps upstream of the start site is more divergent however, and only shares 78% identity between AR-0945 and the Ngo strains used in this study, which is perhaps sufficiently divergent to preclude HGT. Alternative explanations for failed HGT could include: an insufficient amount of donor DNA was provided, DNA methylation using M.CviPI and M.SssI methyltransferases was unsuccessful (28, 29), incompatible DNA uptake sequences (DUSs) (3), or incompatible genomic backgrounds for the expression of these resistance mechanisms; however, follow up studies will be needed to discriminate the underlying cause(s).

Overall, our results expand on prior studies that provide initial insights into illuminating the commensal resistome (17, 21, 22), which is a known source of antibiotic resistance for *Neisseria* pathogens (6, 8-10). We find evidence of constraint on high level macrolide resistance genotypic evolution in AR-0945, with convergence on only two sites in the genome, however different genetic starting places will likely impact evolutionary outcome due to epistatic and additive effects between loci. Thus, future work will not only focus on expanding this approach to other commensal species and therapeutics, but will incorporate intraspecific variation as an additional consideration. Ultimately, this work emphasizes the power of experimental evolution in characterizing the genetic pathways to resistance in commensals species, which will be key to illuminating mutations at risk of transfer across species boundaries and their effects.

## Methods

### Bacterial strains and growth conditions

The bacterial strain *N. elongata* AR Bank #0945 used for this study was obtained from the Centers for Disease Control and Prevention (CDC) and Food and Drug Association’s (FDA) Antibiotic Resistance (AR) Isolate Bank “*Neisseria* species MALDI-TOF Verification panel”. For all experiments, *N. elongata* AR Bank #0945 and its evolved derivatives were revived from trypticase soy broth (TSB) stocks containing 50% glycerol stored at -80°C. Stocks were streaked onto GC agar base (Becton Dickinson Co., Franklin Lakes, NJ, USA) media plates containing 1% Kelloggs solution (38) (GCP-K plates), and were grown for 18-24 hours at 37°C in a 5% CO_2_ atmosphere.

Experimental evolution was conducted by passaging 16 replicate stocks of *N. elongata* AR Bank #0945 in the presence of azithromycin for 20 days or ∼480 generations. A selective gradient of azithromycin was applied to GCB-K plates using Etest strips (bioMérieux, Durham, NC), and each day overnight growth was collected from the entire zone of inhibition (ZOI) and a 1 cm region in the bacterial lawn surrounding the ZOI (Figure 1). Collected cells were suspended in TSB, and plated onto a fresh GCB-K plate with a new Etest strip. MICs each day were determined by reading the lowest concentration that inhibited growth, and reduced susceptibility was determined using the CLSI guidelines for N. gonorrhoeae (breakpoint AZI ≥ 2 μg/mL). MICs were read by at least two independent researchers. Five strains were passaged using the same protocol on media containing no selective agent. Final cell populations were streaked onto GCB-K plates and individual colonies were picked and stocked for further analysis.

### Whole genome sequencing and comparative genomics

Cells from evolved cell lines were lysed by suspending growth from overnight plates in TE buffer (10 mM Tris [pH 8.0], 10 mM EDTA) with 0.5 mg/mL lysozyme and 3 mg/mL proteinase K (Sigma-Aldrich Corp., St. Louis, MO). DNA was purified using the PureLink Genomic DNA Mini kit (Thermo Fisher Corp., Waltham, MA) and treated with RNase A. Sequencing libraries were prepared using the Nextera XT kit (Illumina Corp., San Diego, CA), and uniquely dual-indexed and pooled. Each pool was subsequently sequenced the Illumina MiSeq platform at the Rochester Institute of Technology Genomics Core using V3 600 cycle cartridges (2×300bp). Sequencing quality of each paired-end read library was assessed using FastQC v0.11.9 (39). Trimmomatic v0.39 (40) was used to trim adapter sequences, and remove bases with phred quality score < 15 over a 4 bp sliding window. Reads < 36 bp long, or those missing a mate, were also removed from subsequent analysis. The AR-0945 draft assembly was constructed using SPAdes v.3.14.1 (41) and annotated with prokka v.1.14.5 (42). Trimmed reads were mapped back to the AR-0945 draft assembly using Bowtie2 v.2.2.4 (43) using the “end-to-end” and “very-sensitive” options and Pilon v.1.16 (44) was used to call variant sites. Only single nucleotide polymorphisms and short indels were retained, and any variants called within 100 bp of the end of contigs were removed.

### Transformations

Transformations were conducted by inoculating GCP broth (7.5 g protease peptone 3, 0.5 g soluble starch, 2 g dibasic K2HPO4, 0.5 g monobasic KH2PO4, 2.5 g NaCl, and double-distilled water [ddH2O] to 500 ml; Becton Dickinson Co., Franklin Lakes, NJ) supplemented with 1% IsoVitaleX and 10 mM MgCl (Sigma-Aldrich Corp., St. Louis, MO) with cells to an optical density (OD) of ∼0.5. Cell suspensions were subsequently incubated with 100 ng of gDNA for 4 hours to allow DNA uptake, homologous recombination, and expressions of new alleles. Cell suspensions were then serially diluted to allow for quantification of transformation efficiency and spotted onto GCB-K plates containing 4 μg/mL of azithromycin. Cultures then were incubated overnight and after 18 hours colonies were counted for each reaction and azithromycin resistant transformants were selected by picking single colonies. Transformants were subsequently MIC tested and whole-genome sequenced to nominate inherited derived mutations.

Polymerase chain reaction (PCR) and Sanger sequencing were used to confirm the location and identity of all derived polymorphisms nominated in transformants from genomic sequencing screens. In brief, PCR was conducted in 50-μl volumes using Phusion High-Fidelity DNA polymerase (New England Biolabs, Ipswich, MA). Primers for *rpmH* (F: 5’-CGAAGCTTTCCAAAACGGCT-3’; R: 5’-AAGGTTCGGCCAAAGATTGC-3’) and the *rpsC/rplB* intergenic region (F: 5’-ATCGCTACTTTTAGCAAACCACT-3’; R: 5’-TGCAGAGCATAATGAAGGTGCT-3’) were annealed at 60°C. Reactions were conducted for 35 cycles, with 30 second extensions. Resultant products were cleaned using ExoSAP-IT (Applied Biosystems, Foster City, CA), and sequenced via the Sanger method.

For Ngo transformation experiments, recipient 28Bl and FA1090 cell stocks were obtained from the CDC and Yonatan Grad at the Harvard T.H. Chan School of Public Health respectively. These recipients were provided with genomic DNA from the evolved Nel strains generated within this study both with and without methylation modification. In brief, for modified DNA 500 ng of Nel DNA was incubated with M.CviPI and M.SssI methyltransferases (New England Biolabs, Ipswich, MA) as per the manufacturer’s instructions, and as specified in (29). 100 ng of modified Nel DNA was then provided to 28Bl and FA1090 as transformation substrate, following the aforementioned protocol.

### Growth Curves

Cells were inoculated into GCP broth supplemented 1% Kelloggs solution to an OD_600_ of ∼0.1, using a Genesys 150 spectrophotometer (Thermo Scientific, Waltham, MA). Cell suspensions were subsequently distributed into 96-well plates and incubated at 37°C in a BioTek Synergy H1 microplate reader (BioTek, Winooski, VT). Subsequent OD_600_ measurements were taken every hour for 21 hours, with a 1-minute shake at 180cpm prior to reading. The BioTek Gen5 v.3.05 software was used to interpret OD values. Replicates were completed for each cell line (n=6), and all downstream analyses were performed in R (45).

## Data Availability

All scripts and datasets are available on: https://github.com/wadsworthlab. The draft assembly for *N. elongata* AR Bank #0945 is deposited to GenBank (accession: JAFEUH000000000). Read libraries for the genomics datasets generated in this study can be accessed on the Sequence Read Archive for evolved strains (SAMN17958618-SAMN17958637) and transformants (SAMN18355358-SAMN18355369).

## Acknowledgements

This work was produced by the members of the Fall 2020 Genomics course (BIOL340) at the Rochester Institute of Technology (RIT) – a big thank you to each and every student for their dedication and hard work during an especially trying semester. The authors would like to acknowledge the generous support provided by the RIT College of Science for this study. The authors would also like to thank Narayan H. Wong and the RIT Genomics Core for providing sequencing support.

## Notes

### Competing Interest Statement

The authors have declared no competing interest.

## References

1. Hamilton HL, Dillard JP. 2006. Natural transformation of Neisseria gonorrhoeae: from DNA donation to homologous recombination. Mol Microbiol 59:376–385.

2. Fussenegger M, Rudel T, Barten R, Ryll R, Meyer TF. 1997. Transformation competence and type-4 pilus biogenesis in Neisseria gonorrhoeae – a review. Gene 192:125–134.

3. Frye SA, Nilsen M, Tonjum T, Ambur OH. 2013. Dialects of the DNA Uptake Sequence in Neisseriaceae. PLoS Genet 9:e1003458.

4. Halter R, Pohlner J, Meyer TF. 1989. Mosaic-like organization of IgA protease genes in Neisseria gonorrhoeae generated by horizontal genetic exchange in vivo. EMBO J 8:2737–2744.

5. Feavers IM, Heath AB, Bygraves JA, Maiden MC. 1992. Role of horizontal genetic exchange in the antigenic variation of the class 1 outer membrane protein of Neisseria meningitidis. Mol Microbiol 6:489–495.

6. Spratt BG, Bowler LD, Zhang QY, Zhou J, Smith JM. 1992. Role of interspecies transfer of chromosomal genes in the evolution of penicillin resistance in pathogenic and commensal Neisseria species. J Mol Evol 34:115–125.

7. Zhou J, Spratt BG. 1992. Sequence diversity within the argF, fbp and recA genes of natural isolates of Neisseria meningitidis: interspecies recombination within the argF gene. Mol Microbiol 6:2135–2146.

8. Wadsworth CB, Arnold BJ, Sater MRA, Grad YH. 2018. Azithromycin resistance through interspecific acquisition of an epistasis-dependent efflux pump component and transcriptional regulator in Neisseria gonorrhoeae. mBio 9:e01419–18.

9. Rouquette-Loughlin CE, Reimche JL, Balthazar JT, Dhulipala V, Gernert KM, Kersh EN, Pham CD, Pettus K, Abrams AJ, Trees DL, St Cyr S, Shafer WM. 2018. Mechanistic basis for decreased antimicrobial susceptibility in a clinical isolate of Neisseria gonorrhoeae possessing a mosaic-like mtr efflux pump locus. mBio 9:587–3.

10. Grad YH, Harris SR, Kirkcaldy RD, Green AG, Marks DS, Bentley SD, Trees D, Lipsitch M. 2016. genomic epidemiology of gonococcal resistance to extended- spectrum cephalosporins, macrolides, and fluoroquinolones in the United States, 2000–2013. J Infect Dis 214:1579–1587.

11. Retchless AC, Kretz CB, Chang H-Y, Bazan JA, Abrams AJ, Turner AN, Jenkins LT, Trees DL, Tzeng Y-L, Stephens DS, MacNeil JR, Wang X. 2018. Expansion of a urethritis-associated Neisseria meningitidis clade in the United States with concurrent acquisition of N. gonorrhoeae alleles. BMC Genomics 19:176.

12. Arnold B, Sohail M, Wadsworth C, Corander J, Hanage WP, Sunyaev S, Grad YH. 2020. Fine-scale haplotype structure reveals strong signatures of positive selection in a recombining bacterial pathogen. Mol Biol Evol 37:417–428.

13. Corander J, Connor TR, O’Dwyer CA, Kroll JS, Hanage WP. 2011. Population structure in the Neisseria, and the biological significance of fuzzy species. J Roy Soc Interface 9:1208–1215.

14. Chamorro G, Ibarz-Pavon AB, Kawabata A, León ME, Orrego V, Nagai M, Gabastou JM. 2019. Carriage of Neisseria meningitidis and other Neisseria species among children and young adults in Paraguay. J Med Microbiol 68:1793– 1801.

15. Diallo K, Trotter C, Timbine Y, Tamboura B, Sow SO, Issaka B, Dano ID, Collard J-M, Dieng M, Diallo A, Mihret A, Ali OA, Aseffa A, Quaye SL, Bugri A, Osei I, Gamougam K, Mbainadji L, Daugla DM, Gadzama G, Sambo ZB, Omotara BA, Bennett JS, Rebbetts LS, Watkins ER, Nascimento M, Woukeu A, Manigart O, Borrow R, Stuart JM, Greenwood BM, Maiden MCJ. 2016. Pharyngeal carriage of Neisseria species in the African meningitis belt. J Infect 72:667–677.

16. Unemo M, Dillon J-AR. 2011. Review and international recommendation of methods for typing Neisseria gonorrhoeae isolates and their implications for improved knowledge of gonococcal epidemiology, treatment, and biology. Clin Microbiol Rev 24:447–458.

17. Goytia M, Thompson ST, Jordan SV, King KA. 2021. antimicrobial resistance profiles of human commensal Neisseria species. https://doi.org/10.20944/preprints202103.0522.v1

18. Liu G, Tang CM, Exley RM. 2015. Non-pathogenic Neisseria: members of an abundant, multi-habitat, diverse genus. Microbiol 161:1297–1312.

19. Diallo K, MacLennan J, Harrison OB, Msefula C, Sow SO, Daugla DM, Johnson E, Trotter C, MacLennan CA, Parkhill J, Borrow R, Greenwood BM, Maiden MCJ. 2019. Genomic characterization of novel Neisseria species. Sci Rep 9:13742–11.

20. Dong HV, Pham LQ, Nguyen HT, Nguyen MXB, Nguyen TV, May F, Le GM, Klausner JD. 2019. Decreased cephalosporin susceptibility of oropharyngeal Neisseria species in antibiotic-using men who have sex with men in Hanoi, Vietnam. Clin Infect Dis 70:1169–1175.

21. Fiore MA, Raisman JC, Wong NH, Hudson AO, Wadsworth CB. 2020. Exploration of the Neisseria resistome reveals resistance mechanisms in commensals that may be acquired by N. gonorrhoeae through horizontal gene transfer. Antibiotics 9:656–12.

22. de Block T, Laumen JGE, Van Dijck C, Abdellati S, De Baetselier I, Manoharan-Basil SS, Van den Bossche D, Kenyon C. 2021. WGS of commensal Neisseria reveals acquisition of a new ribosomal protection protein (MsrD) as a possible explanation for high level azithromycin resistance in Belgium. Pathogens 10:384.

23. Jansen G, Barbosa C, Schulenburg H. 2013. Experimental evolution as an efficient tool to dissect adaptive paths to antibiotic resistance. Drug Resist Updates 16:96–107.

24. Gong Z, Lai W, Liu M, Hua Z, Sun Y, Xu Q, Xia Y, Zhao Y, Xie X. 2016. Novel genes related to ceftriaxone resistance found among ceftriaxone-resistant Neisseria gonorrhoeae strains selected in vitro. Antimicrob Agents Ch 60:2043–2051.

25. Laumen JGE, Manoharan-Basil SS, Verhoeven E, Abdellati S, De Baetselier I, Crucitti T, Xavier BB, Chapelle S, Lammens C, Van Dijck C, Malhotra-Kumar S, Kenyon C. 2021. Molecular pathways to high-level azithromycin resistance in Neisseria gonorrhoeae. J Antimicrob Chemother.

26. St Cyr S, Barbee L, Workowski KA, Bachmann LH, Pham C, Schlanger K, Torrone E, Weinstock H, Kersh EN, Thorpe P. 2020. Update to CDC’s treatment guidelines for gonococcal infection, 2020. MMWR Morb Mortal Wkly Rep 69:1911– 1916.

27. Clinical Laboratory Standards Institute. 2019. Performance standards for antimicrobial susceptibility testing, 29th ed. CLSI supplement M100.Clinical and Laboratory Standards Institute, Wayne, PA.

28. So M, Rendón MA. 2019. Tribal warfare: Commensal Neisseria kill pathogen Neisseria gonorrhoeae using its DNA. Microbial Cell 6:544–546.

29. Kim WJ, Higashi D, Goytia M, Rendón MA, Pilligua-Lucas M, Bronnimann M, McLean JA, Duncan J, Trees D, Jerse AE, So M. 2019. Commensal Neisseria kill Neisseria gonorrhoeae through a DNA-dependent mechanism. Cell Host Microbe 26:228–239.e8.

30. Chisholm SA, Dave J, Ison CA. 2010. High-Level azithromycin resistance occurs in Neisseria gonorrhoeae as a result of a single point mutation in the 23S rRNA genes. Antimicrob Agents Ch 54:3812–3816.

31. Ng LK, Martin I, Liu G, Bryden L. 2002. Mutation in 23S rRNA associated with macrolide resistance in Neisseria gonorrhoeae. Antimicrob Agents Ch 46:3020– 3025.

32. Ma KC, Mortimer TD, Duckett MA, Hicks AL, Wheeler NE, Sánchez-Busó L, Grad YH. 2020. Increased power from conditional bacterial genome-wide association identifies macrolide resistance mutations in Neisseria gonorrhoeae. Nat Commun 11:5374–8.

33. Demczuk W, Martin I, Peterson S, Bharat A, Van Domselaar G, Graham M, Lefebvre B, Allen V, Hoang L, Tyrrell G, Horsman G, Wylie J, Haldane D, Archibald C, Wong T, Unemo M, Mulvey MR. 2016. Genomic epidemiology and molecular resistance mechanisms of azithromycin-resistant Neisseria gonorrhoeae in Canada from 1997 to 2014. J Clin Microbiol 54:1304–1313.

34. Hagman KE, Shafer WM. 1995. Transcriptional control of the mtr efflux system of Neisseria gonorrhoeae. J Bacteriol 177:4162–4165.

35. Zarantonelli L, Borthagaray G, Lee EH, Veal W, Shafer WM. 2001. Decreased susceptibility to azithromycin and erythromycin mediated by a novel mtr(R) promoter mutation in Neisseria gonorrhoeae. J Antimicrob Chemoth 47:651–654.

36. Ohneck EA, Zalucki YM, Johnson PJT, Dhulipala V, Golparian D, Unemo M, Jerse AE, Shafer WM. 2011. A novel mechanism of high-level, broad-spectrum antibiotic resistance caused by a single base pair change in Neisseria gonorrhoeae. mBio 2:e00187.–11–e00187–11.

37. Vincent LR, Kerr SR, Tan Y, Tomberg J, Raterman EL, Dunning Hotopp JC, Unemo M, Nicholas RA, Jerse AE. 2018. In vivo-selected compensatory mutations restore the fitness cost of mosaic penA alleles that confer ceftriaxone resistance in Neisseria gonorrhoeae. mBio 9:e01905.#x2013;17–18.

38. Kellogg DS, Peacock WL, Deacon WE, Brown L, Pirkle DI. 1963. Neisseria gonorrhoeae. I. Virulence genetically linked to clonal variation. J Bacteriol 85:1274–1279.

39. Andrews, S. 2010. FastQC: A quality control tool for high throughput sequence data. Available online: http://www.bioinformatics.babraham.ac.uk/projects/fastqc/ (accessed on 28 September 2020).

40. Bolger AM, Lohse M, Usadel B. 2014. Trimmomatic: a flexible trimmer for Illumina sequence data. Bioinformatics 30:2114–2120.

41. Bankevich A, Nurk S, Antipov D, Gurevich AA, Dvorkin M, Kulikov AS, Lesin VM, Nikolenko SI, Pham S, Prjibelski AD, Pyshkin AV, Sirotkin AV, Vyahhi N, Tesler G, Alekseyev MA, Pevzner PA. 2012. SPAdes: A new genome assembly algorithm and its applications to single-cell sequencing. J Comput Biol 19:455–477.

42. Seemann T. 2014. Prokka: rapid prokaryotic genome annotation. Bioinformatics 30:2068–2069.

43. Langmead B, Salzberg SL. 2012. Fast gapped-read alignment with Bowtie 2. Nat Meth 9:357–359.

44. Walker BJ, Abeel T, Shea T, Priest M, Abouelliel A, Sakthikumar S, Cuomo CA, Zeng Q, Wortman J, Young SK, Earl AM. 2014. Pilon: An integrated tool for comprehensive microbial variant detection and genome assembly improvement. PLoS ONE 9:e112963–14.

45. R Core Team. 207. R: A language and environment for statistical computing. R Foundation for Statistical Computing, Vienna, Austria. Available online: https://www.R-project.org/.

